# Distinguishing genetic correlation from causation across 52 diseases and complex traits

**DOI:** 10.1101/205435

**Authors:** Luke J. O’Connor, Alkes L. Price

**Affiliations:** Department of Epidemiology, Harvard T.H. Chan School of Public Health, Boston, MA; Program in Bioinformatics and Integrative Genomics, Harvard Graduate School of Arts and Sciences, Cambridge, MA; Department of Biostatistics, Harvard T.H. Chan School of Public Health, Boston, MA; Program in Medical and Population Genetics, Broad Institute, Cambridge, MA

## Abstract

Mendelian randomization (MR) is widely used to identify causal relationships among heritable traits, but it can be confounded by genetic correlations reflecting shared etiology. We propose a model in which a latent causal variable mediates the genetic correlation between two traits. Under the latent causal variable (LCV) model, trait 1 is *fully genetically causal* for trait 2 if it is perfectly genetically correlated with the latent causal variable, implying that the entire genetic component of trait 1 is causal for trait 2; it is *partially genetically causal* for trait 2 if it has a high genetic correlation with the latent variable, implying that part of the genetic component of trait 1 is causal for trait 2. To quantify the degree of partial genetic causality, we define the *genetic causality proportion* (gcp). We fit this model using mixed fourth moments *E*(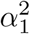*α*_1_*α*_2_) and *E*(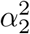*α*_1_*α*_2_) of marginal effect sizes for each trait, exploiting the fact that if trait 1 is causal for trait 2 then SNPs affecting trait 1 (large 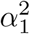) will have correlated effects on trait 2 (large *α*_1_*α*_2_), but not vice versa. We performed simulations under a wide range of genetic architectures and determined that LCV, unlike state-of-the-art MR methods, produced well-calibrated false positive rates and reliable gcp estimates in the presence of genetic correlations and asymmetric genetic architectures; we also determined that LCV is well-powered to detect a causal effect. We applied LCV to GWAS summary statistics for 52 traits (average *N*=331k), identifying partially or fully genetically causal effects (1% FDR) for 59 pairs of traits, including 30 pairs of traits with high gcp estimates (gĉp > 0.6). Results consistent with the published literature included genetically causal effects on myocardial infarction (MI) for LDL, triglycerides and BMI. Novel findings included a genetically causal effect of LDL on bone mineral density, consistent with clinical trials of statins in osteoporosis. These results demonstrate that it is possible to distinguish between genetic correlation and causation using genetic data.

## Introduction

Mendelian Randomization (MR) is widely used to identify potential causal relationships among heritable traits, potentially leading to new disease interventions.^1–12^ Genetic variants that are significantly associated with one trait, the “exposure,” are used as genetic instruments to test for a causal effect on a second trait, the “outcome.” If the exposure does have a causal effect on the outcome, then variants affecting the exposure should affect the outcome proportionally. For example, the MR approach has been used to show that LDL^3, 13^ and triglycerides^4^ (but not HDL^3^) have a causal effect on coronary artery disease (CAD). However, a challenge is that genetic variants can affect both traits pleiotropically, and these pleiotropic effects can induce a genetic correlation, especially when the exposure is polygenic.^2, 11, 12, 14–16^ This challenge can potentially be addressed using curated sets of genetic variants that aim to exclude pleiotropic effects, but curated sets of genetic variants are unavailable for most complex traits (curation is more likely to be successful for molecular traits with simpler genetic architectures). One potential solution has been to apply MR bidirectionally, using genome-wide significant SNPs for each trait in turn.^11,17, 18^ This approach relies on the assumption that if there is no genetically causal relationship, then genome-wide significant SNPs for each trait are equally likely to have correlated effects; however, this assumption can be violated due to differences in trait polygenicity or GWAS sample size.

We introduce a latent causal variable (LCV) model, under which the genetic correlation between two traits is mediated by a latent variable having a causal effect on each trait. A special case of the LCV model is when trait 1 is *fully genetically causal* for trait 2, implying that the entire genetic component of trait 1 is causal for trait 2. More generally, trait 1 may be *partially genetically causal* for trait 2, implying that part of the genetic component of trait 1 is causal for trait 2, and we quantify the degree of partial causality using the *genetic causality proportion* (gcp). In simulations we confirm that LCV, unlike other methods, avoids confounding due to genetic correlations, even in the presence of unequal polygenicity or unequal power between the two traits. Applying LCV to GWAS summary statistics for 52 diseases and complex traits (average *N*=331k), we identify both genetically causal relationships that are consistent with the published literature and novel genetically causal relationships.

## Results

### Overview of methods

The latent causal variable (LCV) model is based on a latent variable *L* that mediates the genetic correlation between the two traits (Figure 1a). Under the LCV model, trait 1 is *fully genetically causal* for trait 2 if it is perfectly genetically correlated with *L*, implying that the entire genetic component of trait 1 is causal for trait 2 (Figure 1b). More generally, trait 1 is *partially genetically causal* for trait 2 if the latent variable has a stronger genetic correlation with trait 1 than with trait 2, implying that part of genetic component of trait 1 is causal for trait 2. In order to quantify the proportion of the genetic component of trait 1 that is causal for trait 2, we define the *genetic causality proportion* (gcp) of trait 1 on trait 2. The gcp ranges between 0 (no genetic causality) and 1 (full genetic causality). A high value of gcp (even if it is not exactly 1) implies that interventions targeting trait 1 are likely to affect trait 2, to the extent that they mimic genetic perturbations to trait 1. (However, we caution that the success of an intervention may depend on its mechanism of action and on its timing relative to disease progression.) An intermediate value of gcp implies that some but not all interventions targeting trait 1 will affect trait 2. For example, a recent study provided evidence consistent with either a fully causal relationship between age at menarche (AAM) and height or a shared hormonal pathway affecting both traits.^11^ If this shared pathway (modeled by our latent variable *L*) has a large causal effect on AAM but a small causal effect on height, then AAM would be partially but not fully genetically causal for height. Indeed, LCV provides evidence for only a partially genetically causal relationship (gĉp = 0.43 (0.10), see below).

**Figure 1:**
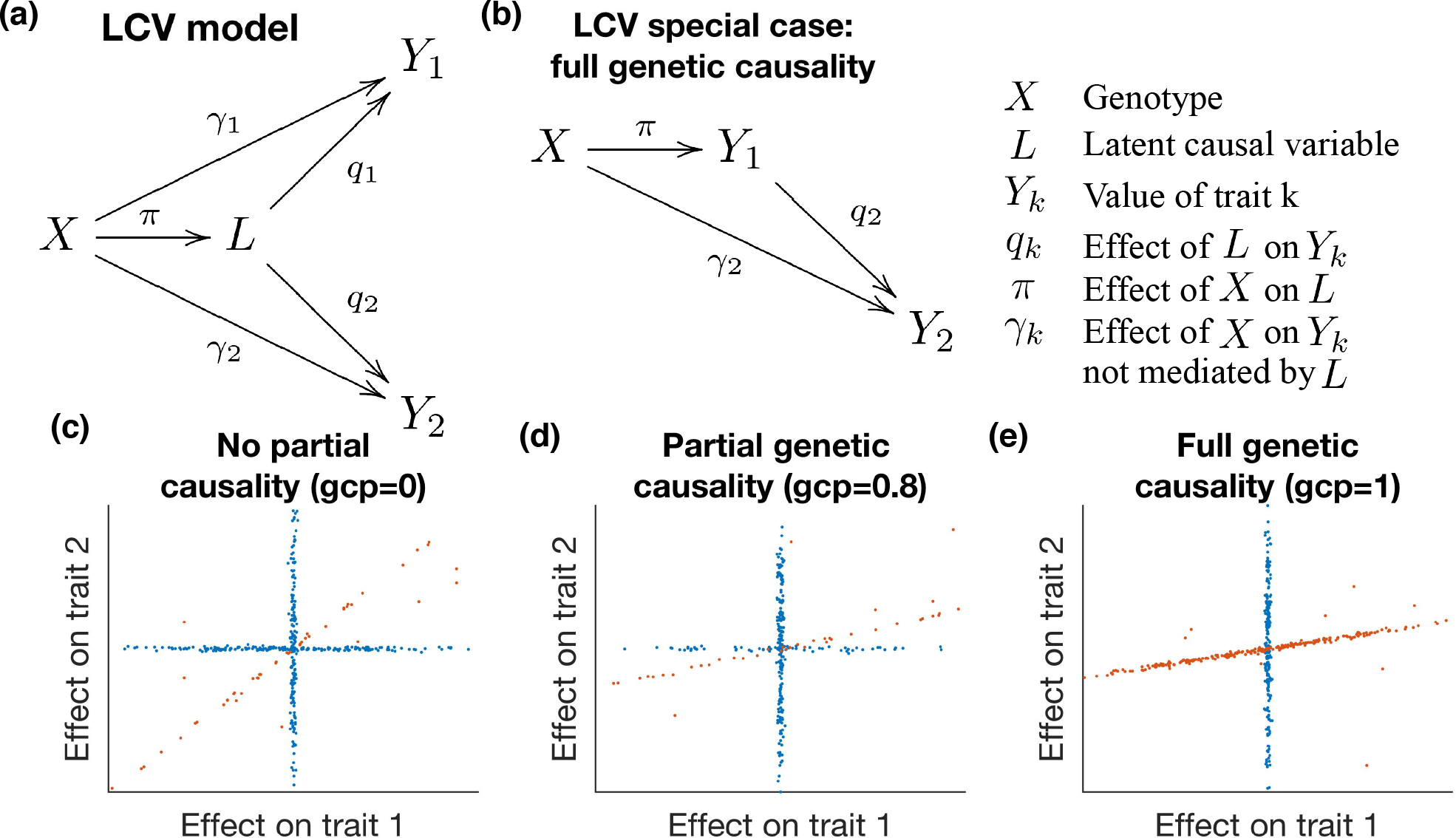
Illustration of the latent causal variable model. We display the relationship between genotypes *X*, latent causal variable *L* and trait values *Y*_1_ and *Y*_2_. (a) Full LCV model. The genetic correlation between traits *Y*_1_ and *Y*_2_ is mediated by a latent variable, *L*, which has normalized effects *q*_1_ and *q*_2_ on each trait. SNPs have random effects *π* on *L* ans random effects *γ*_1_ and *γ*_2_ on each trait. See Table S17 for a list of random variables vs. parameters. (b) When *q*_1_ = 1, *Y*_1_ is perfectly genetically correlated with *L* (so *L* does not need to be shown in the diagram), and we say that *Y*_1_ is fully genetically causal for *Y*_2_. (c) Example genetic architecture of genetically correlated traits with no genetic causality (gcp = 0, i.e. *q*_2_ = *q*_1_ < 1; see equation (2)). Slight noise is added to SNP effects for illustration. Orange SNPs have correlated effects on both traits via *L*, while blue SNPs do not. (d) Example genetic architecture of genetically correlated traits with partial genetic causality (gcp = 0.8, i.e. *q*_2_ < *q*_1_ < 1). The slope of the orange line is determined by the gcp and the genetic correlation. (e) Example genetic architecture of genetically correlated traits with full genetic causality (gcp = 1, i.e. *q*_2_ < *q*_1_ = 1). Under full genetic causality, all SNPs affecting trait 1 also affect trait 2.

In order to test for partial genetic causality and to estimate the gcp, we exploit the fact that if trait 1 is partially genetically causal for trait 2, then most SNPs affecting trait 1 will have proportional effects on trait 2, but not vice versa (Figure 1c-e). Instead of using thresholds to select subsets of SNPs,^11^ we compare the mixed fourth moments *E*(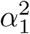*α*_1_*α*_2_) and *E*(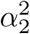*α*_1_*α*_2_) of marginal effect sizes for each trait. The rationale for utilizing these mixed fourth moments is that if trait 1 is causal for trait 2, then SNPs with large effects on trait 1 (i.e. large 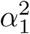) will have proportional effects on trait 2 (large *α*_1_*α*_2_), so that *E*(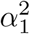*α*_1_*α*_2_) will be large; conversely, SNPs with large effects on trait 2 (large 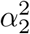) will generally not affect trait 1 (small *α*_1_*α*_2_), so that *E*(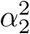*α*_1_*α*_2_) will not be as large. Thus, estimates of the mixed fourth moments can be used to test for partial genetic causality and to estimate the gcp.

LCV assumes that this bivariate distribution is a sum of two independent distributions: (1) a shared genetic component, whose values are proportional for both traits; and (2) a distribution that does not contribute to the genetic correlation (more precisely, a distribution whose density is mirror symmetric across both axes; see Online Methods). We interpret the first distribution as “mediated” effects (corresponding to *π*; see Figure 1a) and the second distribution as “direct” effects (corresponding to *γ*). The LCV model assumption is strictly weaker than the “exclusion restriction” assumption of MR (see Online Methods); in particular, the LCV model permits both correlated pleiotropic effects (mediated by *L*) and uncorrelated pleiotropic effects (not mediated by *L*), while the exclusion restriction assumption permits neither. Under the LCV model, the genetic causality proportion is defined as the number *x* such that:

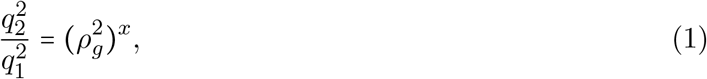

where *q_k_* is the normalized effect of *L* on trait *k* and *ρ_g_* is the genetic correlation^16^ (note that *ρ_g_* = *q*_1_*q*_2_). When the gcp is positive, trait 1 is partially or fully genetically causal for trait 2; when is negative, trait 2 is partially or fully genetically causal for trait 1 (in most instances, we order the two traits so that gcp C 0). We note that partial genetic causality and the gcp can be defined without making LCV (or other) model assumptions (see Online Methods).

In order to estimate the gcp, we utilize the following relationship between the mixed fourth moments of the marginal effect size distribution and the parameters *q*_1_ and *q*_2_:

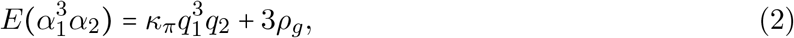

where *π* is the effect of a SNP on *L* and *κ_π_* = *E*(*π*^4^) − 3 is the excess kurtosis of *π* (see Online Methods). This equation implies that if *E*(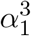*α*_2_)^2^ ≥ *E*(*α*_1_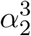)^2^, then 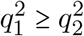.

We calculate statistics *S*(*x*) for each possible value of gcp = *x*, based on equation (2). These statistics utilize estimates of the heritability,^19^ the genetic correlation,^16^ and the cross-trait LD score regression intercept,^16^ in addition to estimates of the mixed fourth moments. We estimate the variance of these statistics using a block jackknife and obtain an approximate likelihood function for the gcp. We compute a posterior mean estimate of gcp (and a posterior standard deviation) using a uniform prior on [−1, 1]. We test the null hypothesis (that gcp = 0) using the statistic *S*(0). Details of the method are provided in the Online Methods section. We have released open source software implementing the LCV method (see URLs).

### Simulations with no LD: comparison with existing methods

To compare the calibration and power LCV with existing causal inference methods, we performed a wide range of null and causal simulations involving simulated summary statistics with no LD. We compared four main methods: LCV, random-effect two-sample MR^5, 9^ (denoted MR), MR-Egger,^7^ and Bidirectional MR^11^ (see Online Methods). We also performed secondary comparisons to two extensions of MR based on the weighted median estimator (MR-WME)^8^ and mode-based estimator (MR-MBE)^10^ (whose performance was roughly similar to MR and to MR-Egger respectively; results of secondary methods are reported in supplementary tables). We applied each method to simulated GWAS summary statistics (*N* = 100k individuals in each of two non-overlapping cohorts; *M* = 50k independent SNPs^20^) for two heritable traits (*h*^2^ = 0.3), generated under the LCV model (simulations under LCV model violations are described below). For each simulation, we display a scatterplot illustrating the bivariate distribution of true SNP effect sizes on the two traits. LCV uses LD score regression^19^ to normalize the summary statistics and cross-trait LD score regression^16^ to estimate the genetic correlation; for simulations with no LD, we use constrained-intercept LD score regression^16^ for both of these steps, resulting in relatively precise estimates of the heritability and the genetic correlation (simulations with LD are described below). A detailed description of all simulations is provided in the Online Methods section, and simulation parameters are described in Table S1.

First, we performed null simulations (gcp = 0) with uncorrelated pleiotropic effects (via *γ*_1_, *γ*_2_; see Figure 1a) and zero genetic correlation. 1% of SNPs were causal for both traits (with independent effect sizes, explaining 20% of heritability for each trait), and 4% of SNPs were causal for each trait exclusively (Figure 2a, Table S2a-d). LCV produced conservative p-values (0.0% false positive rate at *α* = 0.05); our normalization of the test statistic can lead to conservative p-values when the genetic correlation is low (see Online Methods; analyses of real phenotypes are restricted to genetically correlated traits). All three main MR methods produced well-calibrated p-values. Even though the “exclusion restriction” assumption of MR– that there is no pleiotropy– is violated here, these results confirm that uncorrelated pleiotropic effects do not confound random-effect MR at large sample sizes.^21^ We caution that such pleiotropy is known to produce false positives if standard errors are computed using a less conservative fixed-effect approach.^22^ In these simulations, all methods except LCV used the set of approximately 170 SNPs (on average) that were genome-wide significant (*p* < 5 × 10^−8^) for trait 1 (or approximately 330 SNPs that were genome-wide significant for either trait, in the case of Bidirectional MR).

**Figure 2:**
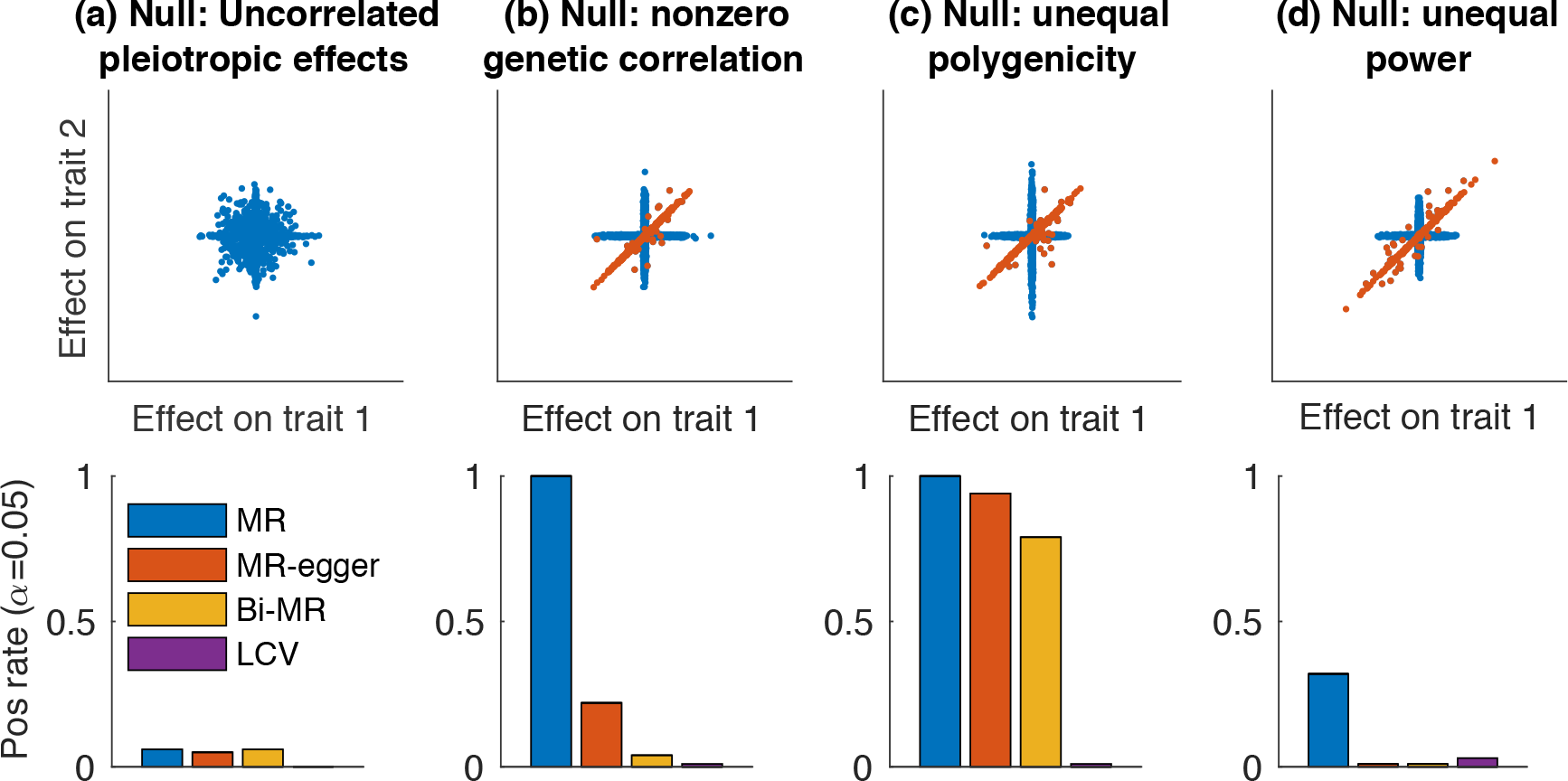
Null simulations with no LD to assess calibration. We compared LCV to three main MR methods (two-sample MR, MR-Egger and Bidirectional MR). We report the positive rate (*α* = 0.05) for a causal (or partially causal) effect. We also display scatterplots illustrating the bivariate distribution of true SNP effect sizes on the two traits. (a) Null simulation (gcp = 0) with uncorrelated pleiotropic effects and zero genetic correlation. (b) Null simulation with nonzero genetic correlation. (c) Null simulation with nonzero genetic correlation and differential polygenicity between the two traits. (d) Null simulation with nonzero genetic correlation and differential power for the two traits. Results for each panel are based on 4000 simulations. Numerical results are reported in Table S2, which also includes comparisons to MR-WME and MR-MBE.

Second, we performed null simulations with a nonzero genetic correlation. 1% of SNPs had causal effects on *L*, and *L* had effects *q*_1_ = *q*_2_ = 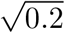on each trait (so that *ρ_g_* = 0.2); 4% of SNPs were causal for each trait exclusively (Figure 2b, Table S2). Even though only ~20% of significant SNPs for the exposure had pleiotropic effects on the outcome, MR and MR-Egger both exhibited severely inflated false positive rates; in contrast, Bidirectional MR and LCV produced well-calibrated p-values. Thus, correlated pleiotropic effects violate the MR exclusion restriction assumption in a manner that leads to false positives. These simulations also violate the MR-Egger assumption that the magnitude of pleiotropic effects on trait 2 are independent of the magnitude of effects on trait 1 (the “InSIDE” assumption),^7^ as SNPs with larger effects on *L* have larger effects on both trait 1 and trait 2 on average, consistent with known limitations.^22^

Third, we performed null simulations with a nonzero genetic correlation and differential polygenicity in the non-shared genetic architecture between the two traits. 1% of SNPs were causal for *L* with effects *q*_1_ = *q*_2_ = 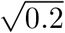on each trait, 2% were causal for trait 1 but not trait 2, and 8% were causal for trait 2 but not trait 1 (Figure 2c, Table S2h-j). Thus, the likelihood that a SNP would be genome-wide significant was higher for causal SNPs affecting trait 1 only than for causal SNPs affecting trait 2 only. We hypothesized that this ascertainment bias would cause Bidirectional MR to incorrectly infer that trait 1 was causal for trait 2. Indeed, Bidirectional MR (as well as other MR methods) exhibited inflated false positive rates, while LCV produced well-calibrated p-values.

Fourth, we performed null simulations with a nonzero genetic correlation and differential power for the two traits, reducing the sample size from 100k to 20k for trait 2. 0.5% of SNPs were causal for *L* with effects *q*_1_ = *q*_2_ = 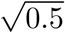 on each trait, and 8% were causal for each trait exclusively (Figure 2d, Table S2k-m). Because per-SNP heritability was higher for shared causal SNPs than for non-shared causal SNPs, shared causal SNPs were more likely to reach genome-wide significance in the smaller trait 1 sample (*N* = 20k); thus, we hypothesized that Bidirectional MR would incorrectly infer that trait 1 was causal for trait 2. Indeed, Bidirectional MR (as well as other MR methods) exhibited inflated false positive rates, while LCV produced well-calibrated p-values.

Next, we performed causal simulations to assess whether LCV is well-powered to detect a causal effect. We consider the relative power of LCV and MR methods to be of secondary importance, given that MR often produces false positives (Figure 2); nonetheless, we report simulation results for the MR methods in addition to LCV, for completeness. We note that LCV had lower power in simulations with LD (see Simulations with LD). First, we chose a set of default parameters: sample size was reduced to *N* = 25k for each trait, 5% of SNPs were causal for trait 1 (the causal trait), there was a (fully) causal effect of size *q*_2_ = 0.2 of trait 1 on trait 2, and an additional 5% of SNPs were causal for trait 2 only (Figure 3a). There were ~15 genome-wide significant SNPs on average, explaining ~2% of *h*^2^. MR and LCV were well-powered to detect a causal effect at *α* = 0.001, while bidirectional MR had lower power and MR-Egger had very low power. Second, we reduced the sample size for trait 1 (Figure 3b, Table S3b-d), finding that LCV had high power while the MR methods had low power, owing to the low number of genome-wide significant SNPs. We caution that LCV produces false positives in simulations with LD with very low sample size, due to noisy heritability estimates, so in practice we restrict our analyses of real traits to data sets with highly significant heritability estimates (see Simulations with LD). Third, we reduced the sample size for trait 2 (Figure 3c, Table S3e-g). LCV and MR had high power, while bidirectional MR and MREgger had low power. Fourth, we reduced the causal effect size of trait 1 on trait 2 (Figure 3d and Table S3h-j). LCV had low power, MR had moderately low power, and other methods had very low power. Fifth, we increased the polygenicity of the causal trait (Figure 3e, Table S3k-m). LCV had high power while the MR methods had low power, again owing to the low number of genome-wide significant SNPs. We also simulated a partially genetically causal relationship (gcp=0.25-0.75), with similar results (Table S3p-r). We compared our gcp estimates in fully causal simulations with our gcp estimates in partially causal simulations, finding that LCV reliably distinguished the two cases, unlike existing methods (Table S3a,p-r). We note that these estimates are biased toward zero due to our use of a mean-zero prior, but we show below that our posterior mean estimates are unbiased in the Bayesian sense (see Simulations with LD). We also note that our estimates of the genetic correlation, which is equal to the size of the causal effect, were unbiased in these simulations (Table S3).

In summary, we determined using simulations with no LD that LCV produced well-calibrated null p-values in the presence of a nonzero genetic correlation, unlike MR and MR-Egger. LCV avoided confounding when polygenicity or power differed between the two traits, unlike Bidirectional MR and other methods (Figure 2). We also determined that LCV was well-powered to detect a true causal effect (Figure 3).

**Figure 3:**
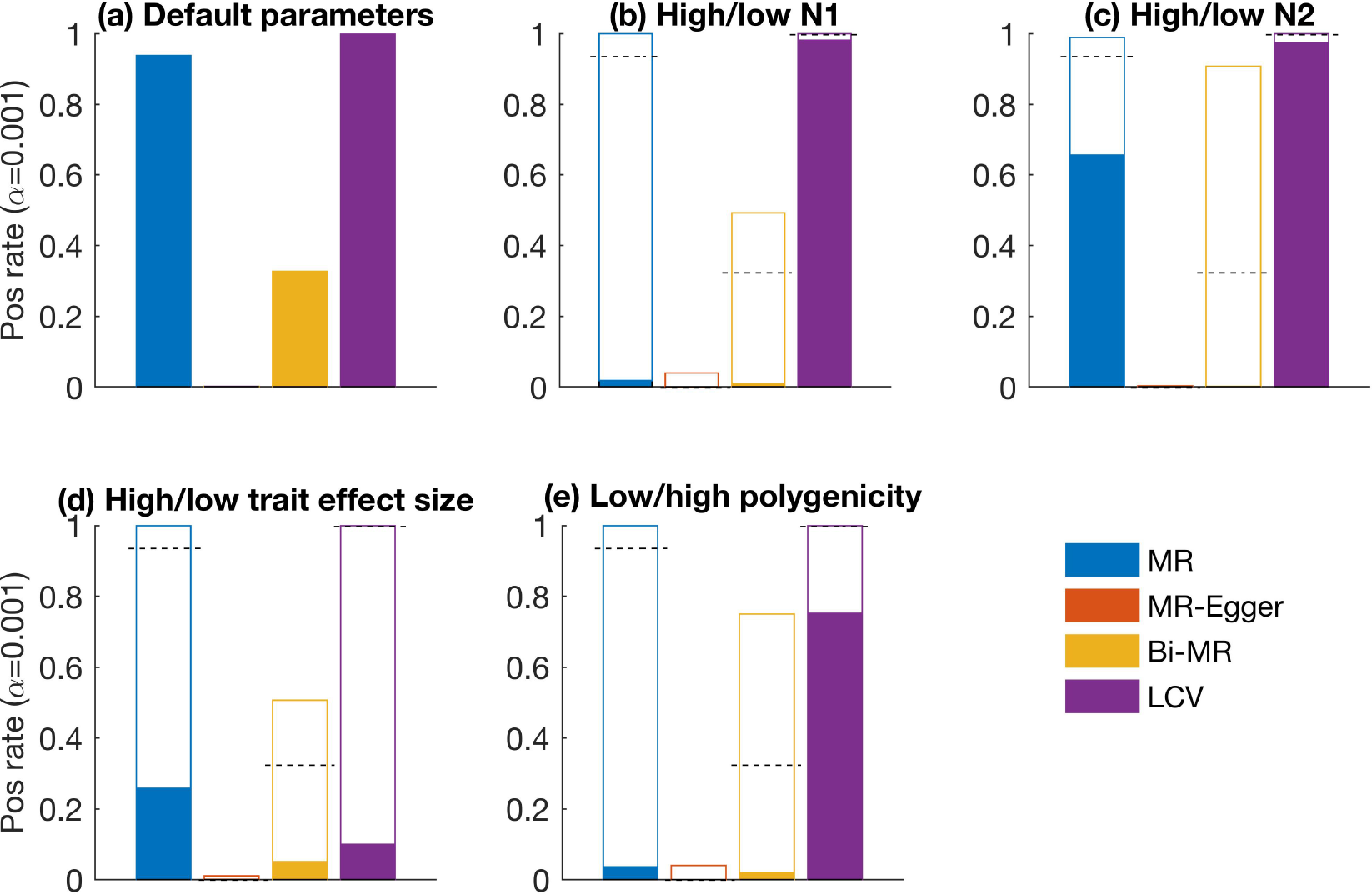
Causal simulations with no LD to assess power. We compared LCV to three main MR methods (two-sample MR, MR-Egger and Bidirectional MR). We report the positive rate (*α* = 0.001) for a causal (or partially causal) effect. (a) Causal simulations with default parameters (results also displayed as dashed lines in panels (b)-(e)). (b) Higher (unfilled) or lower (filled) sample size for trait 1 (the causal trait). (c) Higher (unfilled) or lower (filled) sample size for trait 2. (d) Higher (unfilled) or lower (filled) causal effect size of trait 1 on trait 2. (e) Lower (unfilled) or higher (filled) polygenicity for trait 1 (the causal trait). Results for each panel are based on 1000 simulations. Numerical results are reported in Table S3, which also includes comparisons to MRWME and MR-MBE.

### Simulations with no LD: LCV model violations

In order to investigate potential limitations of our approach, we performed null and causal simulations under genetic architectures that violate LCV model assumptions. As noted above, partial genetic causality is well-defined without making LCV (or other) model assumption (see Online Methods). There are two classes of LCV model violations: *independence violations* and *proportionality violations*. Roughly, independence violations involve a violation of the independence assumption between (1) mediated effects (*π*) and (2) direct effects (γ) while still satisfying a key proportionality condition related to the mixed fourth moments; as a result, independence violations are not expected to cause LCV to produce false positives (see Online Methods). Proportionality violations, on the other hand, violate this proportionality condition and are potentially more problematic. A representative example of an independence violation is a bivariate Gaussian mixture model where one of the mixture components generates imperfectly correlated effect sizes on the two traits. These SNPs underlying this mixture component can be viewed as having both an effect on *L* and also a residual effect on the two traits directly, in violation of the independence assumption. First, we performed null simulations under a Gaussian mixture model with a nonzero genetic correlation. These simulations were similar to the simulations reported in Figure 2b, except that the correlated SNP effect sizes (1% of SNPs) were drawn from a bivariate normal distribution with correlation 0.5 (explaining 20% of heritability for each trait; in Figure 2b, these effects were perfectly correlated). Similar to Figure 2b, LCV and bidirectional MR produced p-values that were well-calibrated, while MR and MR-Egger produced inflated p-values (Figure 4a, Table S4a-d). Second, similar to Figure 2c, we included differential polygenicity between the two traits, finding that differential polygenicity caused all existing methods including bidirectional MR, but not LCV, to produce false positives (Figure 4b, Table S4f-h). Third, similar to Figure 2d, we included differential power between the two traits, again finding that LCV produced well-calibrated p-values while existing methods produced false positives (Figure 4c, Table S4i-k).

**Figure 4:**
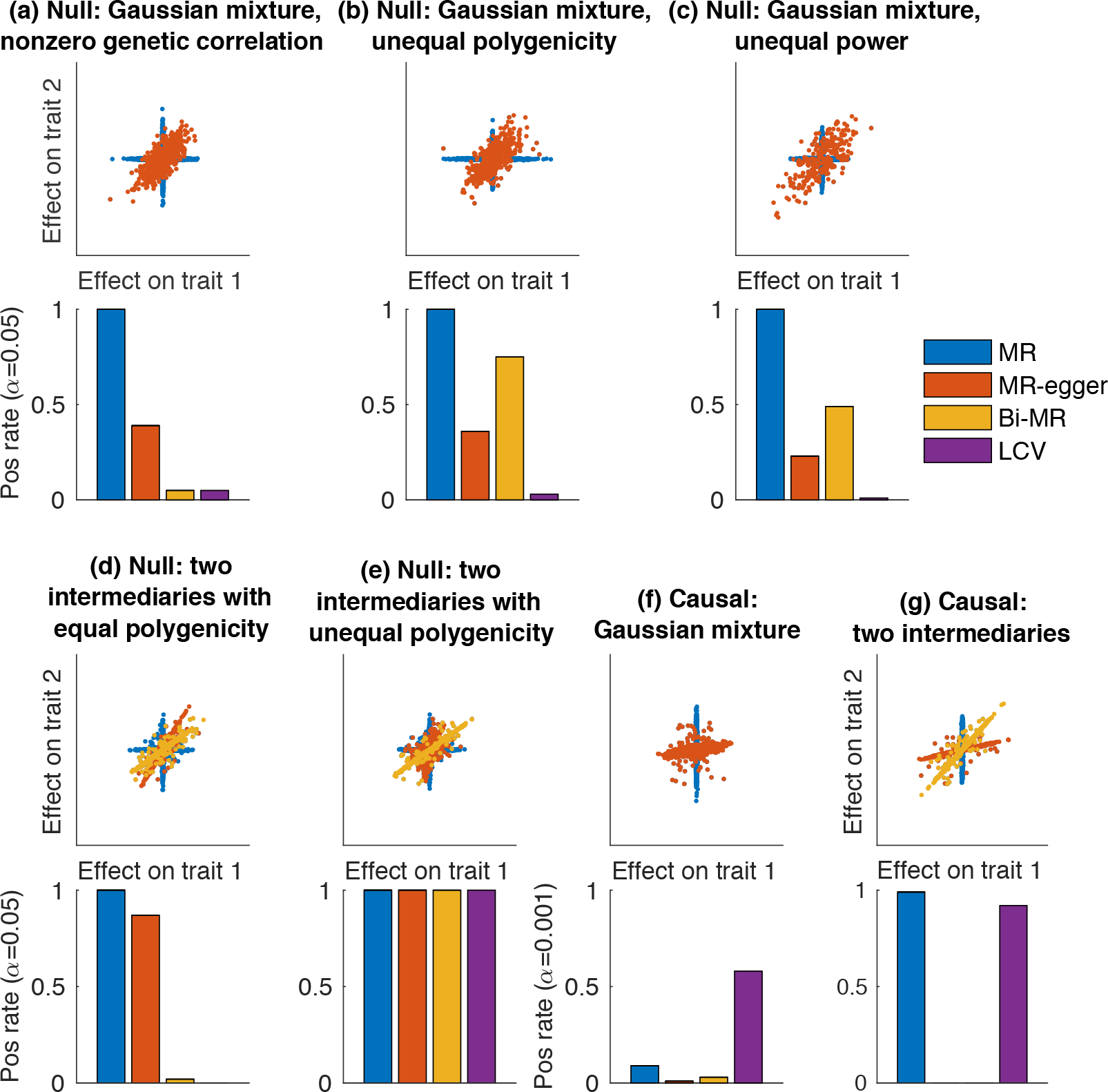
Null and causal simulations with no LD and LCV model violations. We report the positive rate (*α* = 0.05 for null simulations, *α* = 0.001 for causal simulations) for two-sample MR, MR-Egger, Bidirectional MR and LCV. Panels (a)-(c) correspond to Gaussian mixture model extensions of the models in Figure 2b-d. Panels (f) and (g) correspond to causal analogues of the models in panels (a) and (d), respectively. We also display scatterplots illustrating the bivariate distribution of true SNP effect sizes on the two traits. (a) Null simulation with nonzero SNP effects drawn from a mixture of Gaussian distributions; one mixture component has correlated effects on each trait. (b) Null simulation with SNP effects drawn from a mixture of Gaussian distributions, and differential polygenicity between the two traits. (c) Null simulation with SNP effects drawn from a mixture of Gaussian distributions, and unequal power between the two traits. (d) Null simulation with two intermediaries having different effects on each trait. (e) Null simulation with two intermediaries having different effects on each trait and unequal polygenicity for the two intermediaries. (f) Causal simulation with SNP effects drawn from a mixture of Gaussian distributions; all SNPs affecting trait 1 also affect trait 2, but the relative effect sizes were noisy. (g) Causal simulation with an additional genetic confounder (i.e. a second intermediary) mediating part of the genetic correlation. Results for each panel are based on 1000 simulations. Numerical results are reported in Tables S4 and S5, which also includes comparisons to MR-WME and MR-MBE.

A representative example of a proportionality violation is a model in which two intermediaries *L*_1_ and *L*_2_ have different effect sizes on the two traits, and *L*_1_ and *L*_2_ also have unequal polygenicity. First, for comparison purposes, we considered a model with two intermediaries with equal polygenicity; 2% of SNPs were causal for each intermediary, and 4% of SNPs were causal for each trait exclusively. Because this model implies only an independence violation (see Online Methods), we expected that LCV would not produce false positives. Indeed, LCV produced well-calibrated p-values (Figure 4d, Table S5a). Similar to Figure 2b and Figure 4a, Bidirectional MR also produced well-calibrated p-values, while MR and MR-Egger produced false positives. Second, we shifted the polygenicity of the two intermediaries in opposite directions: 1% of SNPs were causal for *L*_1_ and 8% of SNPs were causal for *L*_2_, resulting in a proportionality violation. We expected that LCV would produce false positives, as the intermediary with lower polygenicity would disproportionately affect the mixed fourth moments. Indeed, LCV (as well as other methods) produced false positives, indicating that proportionality violations cause LCV to produce false positives (Figure 4e, Table S5b-d). We investigated the gcp estimates produced by LCV in these simulations, finding that LCV produced low gcp estimates (gĉp ≈ 0.5; Figure S1a). We varied the difference in polygenicity as well as the difference in the relative effect sizes of the two intermediaries, finding that extreme parameter settings (e.g., a 32× difference in polygenicity in conjunction with a 25× difference in the relative effect sizes of *L*_1_ and *L*_2_) were required to cause LCV to produce high gcp estimates (gcp > 0.6; Figure S1a). Thus, proportionality violations of LCV model assumptions can cause LCV (and other methods) to produce false positives, but genetic causality remains the most parsimonious explanation for high gcp estimates.

Finally, we performed (fully) causal simulations under LCV model violations. First, we simulated an independence violation by specifying a Gaussian mixture model where every SNP affecting trait 1 also affected trait 2, but the relative effect sizes were noisy (Figure 4f, Table S4l-n). Sample size and polygenicity were similar to Figure 3a (4× lower sample size than Figure 4a). As expected, LCV had lower power to detect a causal effect than in Figure 3a, although it still had moderately high power. Second, we simulated a proportionality violation by specifying both a causal effect (corresponding to *L*_1_) and an additional genetic confounder (corresponding to *L*_2_) (Figure 4g, Table S5i-k). LCV had lower power to detect a causal effect than in Figure 3a, although it still had high power. We investigated the gcp estimates produced by LCV in these simulations, finding that they were substantially lower than 1 (Table S5i-k and Figure S1b). Therefore, gcp estimates lower than 1 should not be viewed as conclusive evidence against a fully causal effect; an alternative explanation is that model violations cause LCV to underestimate the gcp.

In summary, we determined in null simulations that independence violations do not cause LCV to produce false positives; in addition, these simulations recapitulated the limitations of existing methods that we observed in simulations under the LCV model (Figure 2). Proportionality violations caused LCV (as well as existing methods) to produce false positives; however, extreme values of the simulation parameters were required in order for LCV to produce high gcp estimates. In causal simulations, we determined that both independence and proportionality violations lead to reduced power for LCV (and other methods), as well as downwardly biased gcp estimates for LCV.

### Simulations with LD

Next, we performed simulations with LD; we note that LD can potentially affect the performance of our method, which uses a modified version of LD score regression^16, 19^ to normalize effect size estimates and to estimate genetic correlations. LD was computed using *M* = 596*k* common SNPs in *N* = 145*k* samples of European ancestry from the UK Biobank interim release.^27^ Unlike our simulations with no LD, these simulations also included sample overlap. Because existing methods exhibited major limitations in simulations with no LD (Figure 2 and Figure 4), we restricted these simulations to the LCV method. A detailed description of the simulations is provided in the Online Methods section.

First, we performed null simulations to assess calibration. We chose a set of default parameters similar to Figure 2b and varied each parameter in turn. In particular, similar to Figure 2, these simulations included uncorrelated pleiotropy, genetic correlations, differential polygenicity between the two traits, and differential power between the two traits (Table 1a-f and Table S6a-m). LCV produced approximately well-calibrated or conservative false positive rates. Slight inflation was observed due to noise in our heritability estimates (Table 1a,c-f and Table S6c-m); proper calibration was restored by using constrained-intercept LD score regression^19^ (resulting in more precise heritability estimates) (Table S7a-f). To avoid problems with noisy heritability estimates, we restrict our analyses of real traits to data sets with highly significant heritability estimates (*Z* score for nonzero *h*^2^ = *Z_h_* > 7). We also determined that uncorrected population stratification led to false (Table S9).

**Table 1:** Null and causal simulations with LD. We report the positive rate (*α* = 0.05 and *α* = 0.001) for a causal (or partially causal) effect for LCV, as well as the mean gĉp (gĉp standard error is less than 0.01 in each row). (a) Default parameter values (see text). (b) Zero genetic correlation (*ρ_g_* = 0). (c) Very high genetic correlation (*ρ_g_* = 0.75). (d) Uncorrelated pleiotropic effects. (e) Differential polygenicity (0.2% and 0.8% of SNPs were causal for trait 1 and trait 2, respectively). (f) Differential power (*N*_1_ = 20k and *N*_2_ = 500k). (g) Population stratification. (h) Full genetic causality (gcp = 1). (i) Low trait 1 sample size (*N*_1_ = 20*k*). (j) Weak causal effect (*q*_2_ = *ρ_g_* = 0.1). (k) High trait 1 polygenicity (5% of SNPs causal). (l) Partial genetic causality (gcp = 0.5). Results for each panel are based on 5,000 simulations.

| | | $\rho_g$ | $p_{LCV} < 0.05$ | $p_{LCV} < 0.001$ | Mean gcp |
| --- | --- | --- | --- | --- | --- |
| a | Default parameter values | 0.2 | 0.058 | 0.003 | 0.00 |
| b | Zero genetic correlation | 0 | 0 | 0 | 0.00 |
| c | Very high genetic correlation | 0.8 | 0.058 | 0.002 | -0.00 |
| d | Uncorrelated pleiotropic effects | 0.2 | 0.054 | 0.001 | 0.00 |
| e | Differential polygenicity | 0.2 | 0.062 | 0.002 | 0.01 |
| f | Differential power | 0.2 | 0.063 | 0.004 | -0.01 |
| g | Population stratification | 0.25 | 0.34 | 0.126 | -0.21 |
| h | Causal | 0.2 | 0.97 | 0.94 | 0.76 |
| i | Low $N_1$ | 0.2 | 0.852 | 0.768 | 0.65 |
| j | Weak causal effect | 0.1 | 0.422 | 0.104 | 0.49 |
| k | $Y_1$ more polygenic | 0.2 | 0.155 | 0.004 | 0.28 |
| l | Partially causal | 0.2 | 0.71 | 0.35 | 0.56 |

Second, we performed causal simulations to assess power. We chose a set of default parameters similar to our null simulations, finding that LCV was well-powered (Table 1h), although its power was lower than in simulations with no LD (Figure 3a). We varied each parameter in turn, finding that power was reduced when we reduced the sample size, increased the polygenicity of the causal trait, reduced the causal effect size, or simulated a partially causal rather than fully causal genetic architecture (Table 1i-l and Table S6u-bb), similar to simulations with no LD (Figure 3b-f). These simulations indicate that LCV is well-powered to detect a causal effect for large GWAS under most realistic parameter settings, although its power does depend on genetic parameters that are difficult to predict.

Third, to assess the unbiasedness of gcp posterior mean (and variance) estimates, we performed simulations in which the true value of gcp was drawn uniformly from [−1, 1] (corresponding to the prior that LCV uses to compute its posterior mean estimates, see Online Methods). We expected posterior-mean estimates to be unbiased in the Bayesian sense that *E*(gcp|gĉp) = gĉp (which differs from the usual definition of unbiasedness, that *E*(gĉp|) = gcp).^26^ Thus, we binned these simulations by gĉp and plotted the mean value of gcp within each bin (Figure S2). We determined that mean gcp within each bin was concordant with gĉp. In addition, the root mean squared error was 0.15, approximately consistent with the root mean posterior variance estimate of 0.13 (Table S8).

In summary, we confirmed using simulations with LD that LCV produces well-calibrated false positive rates under a wide range of realistic genetic architectures; some p-value inflation was observed when heritability estimates were noisy, but false positives can be avoided in analyses of real traits by restricting to traits with highly significant heritability (*Z_h_* > 7). We also confirmed that LCV is well-powered to detect a causal effect under a wide range of realistic genetic architectures, and produces unbiased posterior mean estimates of the gcp.

### Application to real phenotypes

We applied LCV and the MR methods to GWAS summary statistics for 52 diseases and complex traits, including summary statistics for 37 UK Biobank traits^27,28^ computed using BOLT-LMM^29^ (average *N*=429k) and 15 other traits (average *N*=54k) (see Table S10 and Online Methods). The 52 traits were selected based on the significance of their heritability estimates (*Z_h_* > 7), and traits with very high genetic correlations (|*ρ_g_*| > 0.9) were pruned, retaining the trait with higher *Z_h_*. As in previous work, we excluded the MHC region from all analyses, due to its unusually large effect sizes and long-range LD patterns.^19^

We applied LCV to the 429 trait pairs (32% of all trait pairs) with a nominally significant genetic correlation (*p* < 0.05), detecting significant evidence of full or partial genetic causality for 59 trait pairs (FDR < 1%), including 30 trait pairs with gĉp > 0.6. We primarily focus on trait pairs with high gcp estimates, which are of greatest biological interest (and extremely unlikely to be false positives; see Simulations with no LD: LCV model violations and Figure S1a). Results for selected trait pairs are displayed in Figure 5, results for the 30 trait pairs with gĉp > 0.6 are reported in Table 2, results for all 59 significant trait pairs are reported in Table S11, and results for all 429 genetically correlated trait pairs are reported in Table S12.

**Figure 5:**
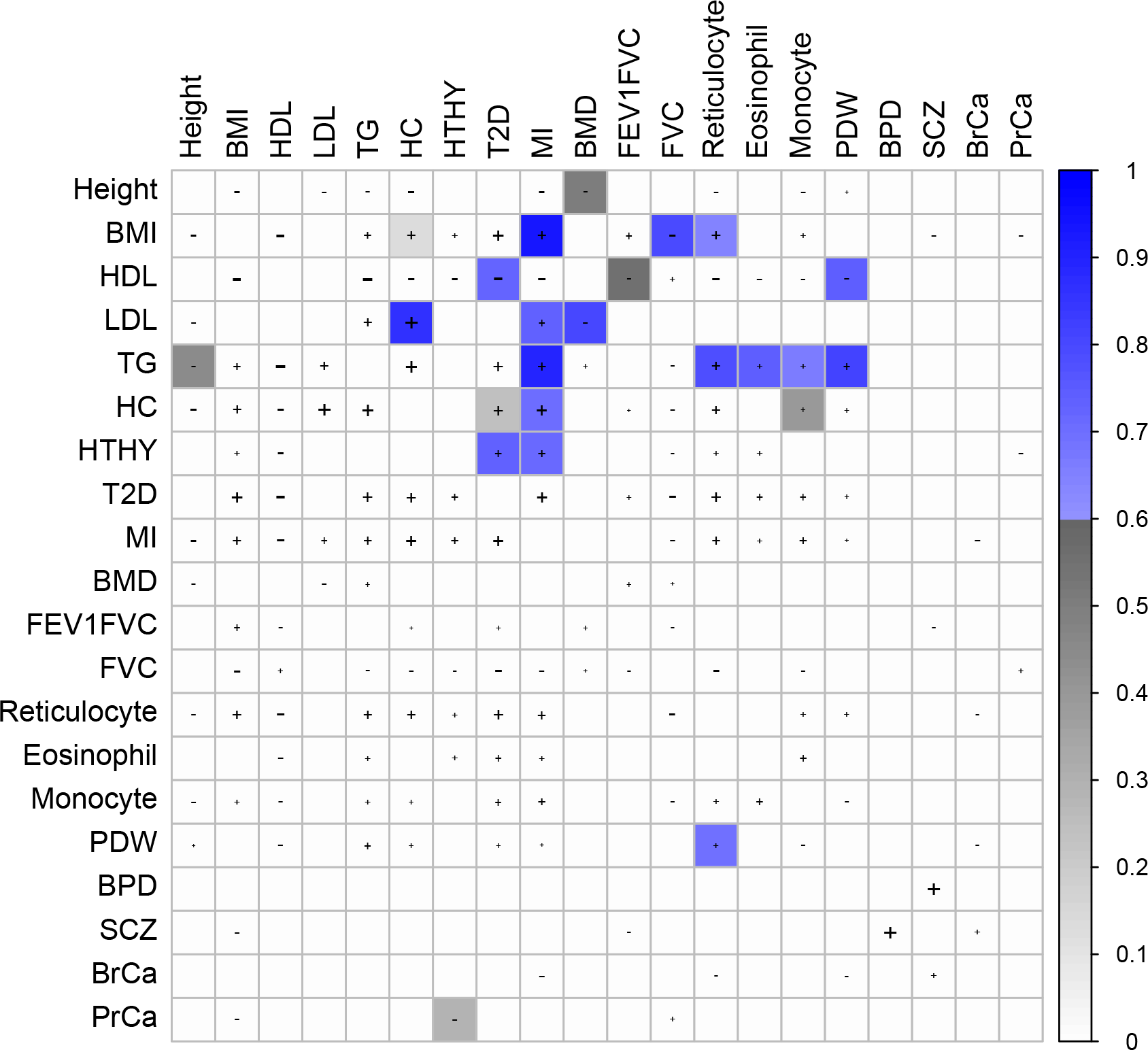
Partially or fully genetically causal relationships between selected complex traits. Shaded squares indicate significant evidence for a causal or partially causal effect of the row trait on the column trait (FDR < 1%). Color scale indicates posterior mean gĉp for the effect of the row trait on the column trait; blue color indicates gĉp > 0.6, grey color indicates gĉp < 0.6. “+” or “-” signs indicate trait pairs with a nominally significant (positive or negative) genetic correlation (*p* < .05), and the size of the “+” or “-” size is proportional to the genetic correlation. Results for the 30 trait pairs with gĉp > 0.6 are reported in Table 2, results for all 59 significant trait pairs are reported in Table S11, and results for all 429 genetically correlated trait pairs are reported in Table S12. HTHY: hypothyroidism. FG: fasting glucose. PDW: platelet distribution width. BPD: bipolar disorder. SCZ: schizophrenia. BrCa: breast cancer: PrCa: prostate cancer.

**Table 2:** Fully or partially genetically causal relationships between complex traits. We report all significant trait pairs (1% FDR) with high gcp estimates (gĉp > 0.6). *p*_LCV_ is the p-value for the null hypothesis of no partial genetic causality; *ρ̂_g_* is the estimated genetic correlation, with standard error; gĉp is the posterior mean estimated genetic causality proportion, with posterior standard error. We provide references for all MR studies supporting causal relationships between these traits that we are currently aware of. Results for all 59 significant trait pairs are reported in Table S11, and results for all 429 genetically correlated trait pairs are reported in Table S12.

| Trait 1 | Trait 2 | $p_{LCV}$ | $\hat{\rho}_g$ (std err) | $\hat{g}cp$(std err) | MR ref |
| --- | --- | --- | --- | --- | --- |
| Triglycerides | Hypertension | $5 \times 10^{-39}$ | 0.25 (0.04) | 0.95 (0.04) | |
| BMI | Heart attack | $3 \times 10^{-9}$ | 0.34 (0.09) | 0.94 (0.11) | 32,38 |
| Triglycerides | Heart attack | $8 \times 10^{-32}$ | 0.30 (0.06) | 0.90 (0.08) | 4 |
| Triglycerides | BP - systolic | $6 \times 10^{-41}$ | 0.13 (0.03) | 0.89 (0.08) | |
| HDL | Hypertension | $6 \times 10^{-22}$ | -0.29 (0.06) | 0.87 (0.09) | |
| LDL | High cholesterol | $8 \times 10^{-7}$ | 0.77 (0.07) | 0.86 (0.11) | |
| Triglycerides | Mean cell volume | $10 \times 10^{-19}$ | -0.20 (0.04) | 0.86 (0.11) | |
| Triglycerides | BP - diastolic | $5 \times 10^{-39}$ | 0.11 (0.04) | 0.86 (0.10) | |
| Platelet volume | Platelet count | $6 \times 10^{-10}$ | -0.66 (0.03) | 0.84 (0.10) | |
| BMI | Hypertension | $2 \times 10^{-16}$ | 0.38 (0.03) | 0.83 (0.11) | 11,38 |
| Triglycerides | Platelet dist width | $5 \times 10^{-17}$ | 0.19 (0.04) | 0.81 (0.13) | |
| LDL | BMD | $4 \times 10^{-34}$ | -0.12 (0.05) | 0.80 (0.12) | |
| BMI | FVC | $4 \times 10^{-13}$ | -0.22 (0.03) | 0.79 (0.17) | 72 |
| Triglycerides | Reticulocyte count | $2 \times 10^{-10}$ | 0.33 (0.05) | 0.79 (0.14) | |
| Triglycerides | Eosinophil count | $3 \times 10^{-17}$ | 0.14 (0.05) | 0.75 (0.16) | |
| Balding | Num children - male | $2 \times 10^{-30}$ | -0.16 (0.05) | 0.75 (0.13) | |
| HDL | Platelet dist width | $8 \times 10^{-17}$ | -0.14 (0.04) | 0.75 (0.16) | |
| RBC dist width | Type 2 Diabetes | $3 \times 10^{-4}$ | 0.11 (0.03) | 0.73 (0.19) | |
| LDL | Heart attack | $2 \times 10^{-31}$ | 0.17 (0.08) | 0.73 (0.13) | 3,13 |
| Platelet dist width | Platelet count | $1 \times 10^{-7}$ | -0.47 (0.04) | 0.73 (0.15) | |
| Hypothyroidism | Type 2 Diabetes | $2 \times 10^{-4}$ | 0.22 (0.05) | 0.73 (0.29) | |
| HDL | Type 2 Diabetes | $2 \times 10^{-7}$ | -0.40 (0.06) | 0.72 (0.17) | |
| Hypothyroidism | Heart attack | $6 \times 10^{-12}$ | 0.26 (0.05) | 0.72 (0.16) | |
| High cholesterol | Heart attack | $2 \times 10^{-4}$ | 0.52 (0.12) | 0.71 (0.19) | |
| HDL | BP - diastolic | $4 \times 10^{-17}$ | -0.12 (0.06) | 0.70 (0.18) | |
| Platelet dist width | Reticulocyte count | $1 \times 10^{-7}$ | 0.13 (0.04) | 0.69 (0.20) | |
| LDL | College | $1 \times 10^{-10}$ | -0.13 (0.05) | 0.68 (0.30) | |
| Triglycerides | Monocyte count | $1 \times 10^{-4}$ | 0.14 (0.04) | 0.67 (0.21) | |
| Type 2 Diabetes | Ulcerative Colitis | $2 \times 10^{-5}$ | -0.14 (0.07) | 0.65 (0.23) | |
| BMI | Reticulocyte count | $4 \times 10^{-5}$ | 0.39 (0.03) | 0.64 (0.25) | |

Myocardial infarction (MI) had a nominally significant genetic correlation with 31 other traits, of which six had significant evidence (FDR < 1%) for a fully or partially genetically causal effect on MI (Table 2); there was no evidence for a genetically causal effect of MI on any other trait. Consistent with previous studies, these traits included LDL,^3, 13^ triglycerides^4^ and BMI,^30^ but not HDL.^3^ The effect of BMI was also consistent with prior MR studies,^30–33^ although these studies did not attempt to account for pleiotropic effects (also see ref. 34, which detected no effect). There was also evidence for a genetically causal effect of high cholesterol, which was unsurprising (due to the high genetic correlation with LDL) but noteworthy because of its strong genetic correlation with MI, compared with LDL and triglycerides. The result for HDL and MI did not pass our significance threshold (FDR < 1%), but was nominally significant (*p* = 0.02, Table S12); we residualized HDL summary statistics on summary statistics for three established causal risk factors (LDL, BMI and triglycerides), determining that residualized HDL remained genetically correlated with MI but showed no evidence of genetic causality (*p* = 0.8). On the other hand, most of the six traits with significant causal effects on MI remained significant after residualizing on the established causal risk factors (Table S13). However, we caution that applying LCV to summary statistics that have been residualized on another complex trait, or computed using a complex trait as a covariate, can lead to false-positive or false-negative results due to collider bias;^35, 36^ therefore, we do not recommend applying LCV to residualized summary statistics as a primary analysis. We confirmed that self-reported MI in UK Biobank was highly genetically correlated with CAD in CARDIoGRAM consortium data^39^ (*ρ̂_g_* = 1.34(0.25); not significantly different from 1).

We also detected evidence for a fully or partially genetically causal effect of hypothyroidism on MI (Table 2). Although hypothyroidism is not as well-established a cardiovascular risk factor as high LDL, its genetic correlation with MI is comparable (Table 2), and this effect is mechanistically plausible.^40, 41^ While this result was robust in the conditional analysis (Table S13), and there was no strong evidence for a genetically causal effect of hypothyroidism on lipid traits (Table S12), it is possible that this effect is mediated by lipid traits. A recent MR study of thyroid hormone levels, at ~20× lower sample size than the present study, provided evidence for a genetically causal effect on LDL but not CAD.^42^ On the other hand, clinical trials have demonstrated that treatment of subclinical hypothyroidism using levothyroxine leads to improvement in several cardiovascular risk factors.^43–47^ We also detected evidence for a fully or partially genetically causal effect of hypothyroidism on T2D (Table S11), consistent with a longitudinal association between subclinical hypothyroidism and diabetes incidence,^48^ as well as an effect of thyroid hormone withdrawal on glucose disposal in athyreotic patients.^49^

We identified four traits with evidence for a fully or partially genetically causal effect on hypertension (Table 2), which is genetically correlated with MI (*ρ̂_g_* = 0.49(0.10)). These included genetically causal effects of BMI, consistent with the published literature,^11, 38^ as well as triglycerides and HDL. The genetically causal effect of HDL indicates that there exist major metabolic pathways affecting hypertension with little or no corresponding effect on MI. The positive partially genetically causal effect of reticulocyte count, which had a low gcp estimate (gĉp = 0.41(0.13)), is likely related to the substantial genetic correlation of reticulocyte count with triglycerides (*ρ̂_g_* = 0.33(0.05)) and BMI (*ρ̂_g_* = 0.39(0.03)).

We detected evidence for a (negative) genetically causal effect of LDL on bone mineral density (BMD; Table 2). A meta-analysis of seven randomized clinical trials reported that statin administration increased bone mineral density, although these clinical results have generally been interpreted as evidence of a shared pathway affecting LDL and BMD.^50^ Moreover, familial defective apolipoprotein B leads to high LDL cholesterol and low bone mineral density.^51^ To further validate this result, we performed two-sample MR using 8 SNPs that were previously used to show that LDL affects CAD (in ref. 3; see Online Methods), finding modest evidence for a (negative) causal effect (*p* = 0.04). Because there is a clear mechanistic hypothesis linking each of these variants to LDL directly, this analysis provides separate evidence for a genetically causal effect (LCV does not prioritize variants that are more likely to satisfy instrumental variable assumptions). We also detected a partially genetically causal effect of height on BMD, with a lower gcp estimate (Table 2).

We detected evidence for a fully or partially genetically causal effect of triglycerides on five cell blood traits: mean cell volume, platelet distribution width, reticulocyte count, eosinophil count and monocyte count (Table 2). These results highlight the pervasive effects of metabolic pathways, which can induce genetic correlations with cardiovascular phenotypes. For example, shared metabolic pathways may explain the high genetic correlation of reticulocyte count with MI (*ρ̂_g_*=0.31(0.06)) and hypertension (*ρ̂_g_* = 0.27(0.04)).

Although we have focused primarily on the 30 trait pairs with high gcp estimates (gĉp > 0.6), approximately half of significant trait pairs had lower gcp estimates. Given that there is lower power to detect trait pairs with low gcp values (Table S3a,p-r), it is likely that partial genetic causality with gcp < 0.6 is more common than full or nearly-full genetic causality with gcp > 0.6. Trait pairs with low gcp estimates can suggest plausible biological hypotheses. For example, we identified a partially genetically causal effect of age at menarche (AAM) on height (gĉp = 0.43(0.10), Table S11), suggesting that these traits are influenced by a shared hormonal pathway that is more strongly correlated with AAM than with height, as recently hypothesized.^11^

A recent study reported genetic correlations between various complex traits and number of children in males and females.^52^ We identified only one trait (balding in males) with a fully or partially causal effect on number of children (negative effect on number of children in males; Table 2). For college education, which has a strong negative genetic correlation with number of children (*ρ̂_g_* = −0.31(0.07) and −0.26(0.06) in males and females respectively), we obtained low gcp with low standard errors (gĉp = 0.00(0.09) and gĉp = 0.04(0.21) respectively, Table S12), providing evidence against causality. Thus, a genetic correlation with number of children does not imply a causal effect. This result does not contradict the conclusion of reference 52 that complex traits are under natural selection, as natural selection produces a change in the mean value of a trait even if the trait is non-causally correlated with fitness.^53^

It has been reported that polygenic autism risk is positively genetically correlated with educational attainment^16^ (and cognitive ability,^54^ a highly genetically correlated trait^57^), possibly consistent with the hypothesis that common autism risk variants are maintained in the population by balancing selection.^55, 56^ If balancing selection involving a trait related to educational attainment explained a majority of autism risk, we would expect that most common variants affecting autism risk would also affect educational attainment, leading to a partially genetically causal effect of autism on educational attainment. However, we detected evidence against such an effect (gĉp = 0.13(0.13), *ρ̂_g_* = 0.23(0.07); Table S12); thus, balancing selection acting on educational attainment or a closely related trait is unlikely to explain the high prevalence of autism.

Several causal or plausibly causal relationships were not identified by LCV (Table S14). We note that non-significant LCV p-values do not constitute evidence against a causal effect, (confidently low gcp estimates do constitute evidence against a causal effect, but LCV did not produce confidently low gcp estimates for most trait pairs discussed below; Table S14). First, LCV did not identify a causal effect of BMI on T2D, due to two outlier loci that do not support a causal effect. After applying an outlier removal procedure to remove these loci (see Online Methods), LCV provides convincing evidence for a fully or partially genetically causal effect (*p* = 9×10^−6^). Pleiotropic outlier loci can cause LCV to produce false negatives (but not false positives); however, this phenomenon appears to be uncommon (Table S15), and we generally do not recommend removing outlier loci because they may contain valuable information. Second, LCV did not identify a causal effect of asthma on pulmonary function (FVC or FEV1/FVC). A possible explanation is diagnosis bias: if individuals with low pulmonary function (for reasons unrelated to asthma) are more likely to be diagnosed with asthma, then this bias would mask the causal effect of asthma on pulmonary function. Third, LCV did not identify a causal effect of smoking status on pulmonary function or MI. A possible explanation is that many SNPs affect smoking status only indirectly, with a primary effect on smoking heaviness or deepness of inhalation.^59^ Such SNPs would have much larger effects on cardiopulmonary traits than would be expected based on their effect on smoking status. This type of pleiotropy causes LCV to have lower power (Figure 4f). Fourth, LCV did not identify a causal effect of anorexia on BMI. A possible explanation is the high polygenicity of anorexia, as LCV has lower power when the polygenicity of the causal trait is high (Figure S3e). We note that for most of the trait pairs described above, Bidirectional MR also did not detect a causal effect (Table S14).

Although none of the significant trait pairs with high gcp estimates (Table 2 are obvious false positives, we assessed the likelihood of false positives due to LCV model violations using an auxiliary test for partial genetic causality that does not rely on LCV model assumptions; this test directly estimates the correlated mixture component of the bivariate distribution of SNP effect sizes and compares the proportion of heritability explained by this correlated component for each trait. Simulations indicate that this test is robust to both types of LCV model violations (Table S4 and S5), though less powerful and prone to false positives due to unequal power, hence not recommended for broad use (Tables S2 and S3); see Supplementary Note for a description of the method and its performance in simulations. We applied the auxiliary test to the 30 trait pairs with high gcp estimates, finding that the estimated direction of effect was concordant with LCV for 30/30 trait pairs. While the auxiliary test replicated the LCV result at a nominal significance level (single-tailed *p* < 0.05) for only 17/30 trait pairs, the fraction 17/30 is expected to be an underestimate of the true positive rate, due to limited power. Indeed, when we applied the auxiliary test to the remaining 394 trait pairs, it produced positive results at the corresponding significance level (two-tailed *p* < 0.10) for only 41/394 trait pairs (39 expected under the null; includes 7/29 trait pairs that LCV reported as significant with gĉp < 0.6). This analysis confirms that the 30 trait pairs reported in Table 2 are extremely unlikely to be false positives.

In order to evaluate whether the limitations of MR observed in simulations (Figure 2) are also observed in analyses of real traits, we applied MR, MR-Egger and Bidirectional MR to all 429 genetically correlated trait pairs (Table S12). MR reported significant causal relationships (1% FDR) for 271/429 trait pairs, including 155 pairs of traits for which each trait was reported to be causal for the other trait. This confirms that MR frequently produces false positives in the presence of a genetic correlation, as predicted by our simulations (Figure 2). In contrast, LCV reported a significant partially or fully genetically causal relationship for only 59 trait pairs (Table S11), and never reported a causal effect in both directions. Similarly, Bidirectional MR reported a significant causal relationship for only 45 trait pairs (including 17 pairs of traits that overlapped with LCV), and never reported a causal effect in both directions. We provide a comparison between the results of LCV and Bidirectional MR in Table S16.

## Discussion

We have introduced a latent causal variable (LCV) model to identify causal relationships among genetically correlated pairs of complex traits. We applied LCV to 52 traits, finding that many trait pairs do exhibit partially or fully genetically causal relationships. Our results included several novel findings, including a genetically causal effect of LDL on bone mineral density (BMD) which suggests that lowering LDL may have additional benefits besides reducing the risk of cardiovascular disease.

Our method represents an advance for two main reasons. First, LCV reliably distinguishes between genetic correlation and full or partial genetic causation. Unlike existing MR methods, LCV provided well-calibrated false positive rates in null simulations with a nonzero genetic correlation, even in simulations with differential polygenicity or differential power between the two traits. Thus, positive findings using LCV are more likely to reflect true causal effects. Second, we define and estimate the genetic causality proportion (gcp) to quantify the degree of causality. This parameter, which provides information orthogonal to the genetic correlation or the causal effect size, enables a non-dichotomous description of the causal architecture. Even when both MR and LCV provide significant p-values, the p-value alone is consistent with either fully causal or partially causal genetic architectures, limiting its interpretability; our gcp estimates appropriately describe the range of likely hypotheses.

This study has several limitations. First, the LCV model includes only a single intermediary and can be confounded in the presence of multiple intermediaries, in particular when the intermediaries have differential polygenicity (Figure 4e). Indeed, some trait pairs with low gcp estimates are potentially consistent with such a phenomenon (Table S11). However, the 30 trait pairs with gĉp > 0.6 reported in Table 2 are unlikely to be false positives, both because our simulations show model violations generally do not lead to high gcp estimates (see Figure S1a) and because the estimated direction of effect for all 30 trait pairs was concordant in an analysis using an auxiliary method that is robust to violations of LCV model assumptions (see Supplementary Note and Table S11). Second, because LCV models only two traits at a time, it cannot be used to identify conditional effects given observed confounders.^4, 60^ This approach was used, for example, to show that triglycerides affect coronary artery disease risk conditional on LDL.^4^ However, it is less essential for LCV to model observed genetic confounders, since LCV explicitly models a latent genetic confounder. Third, LCV can be susceptible to false negatives due to outlier loci, bias in disease diagnosis, strong pleiotropic effects, or a highly polygenic causal trait (Table S14). However, LCV is well-powered to detect a causal effect in most simulations (Figure 3), and it detects many established causal relationships among real traits with very high statistical significance (Table 2). Fourth, LCV is not currently applicable to traits with small sample size and/or heritability, due to low power as well as incorrect calibration. However, GWAS summary statistics at large sample sizes have become publicly available for increasing numbers of diseases and traits, including UK Biobank traits.^29^ Fifth, the LCV model can be confounded confounded by shared population stratification, so it is critical for association statistics to be corrected for stratification. Sixth, while many trait pairs have high gcp estimates (gĉp > 0.6), it is not clear whether most of these trait pairs reflect fully or partially genetically causal relationships. A gcp of 1 and a gcp of ~0.6 would be interpreted differently, as a gcp of ~0.6 suggests that only some interventions on trait 1 will modify trait 2, depending on their mechanism of action. This type of uncertainty can be reduced at higher sample size, but not eliminated entirely. Seventh, even full genetic causality must be interpreted with caution before designing disease interventions, as interventions may fail to mimic genetic perturbations. For example, factors affecting a developmental phenotype such as height might need to be modified at the correct developmental time point in order to have any effect; this limitation broadly applies to all methods for inferring causality using genetic data. Eighth, LCV does not model LD explicitly (unlike cross-trait LD score regression^16^), and consequently it models the marginal, rather than the causal, effect size distribution. Modeling the causal effect size distribution while explicitly accounting for LD would enable LCV analyses to be conditioned on various functional annotations, enabling models involving different shared genetic components such as SNPs linked to gene regulation in different cell types. Ninth, power might also be increased by including rare and low-frequency variants; even though these SNPs explain less complex trait heritability than common SNPs,^20, 61^ they may contribute significantly to power if the genetic architecture among these SNPs is more sparse than among common SNPs. Tenth, we cannot infer whether inferred causal effects are linear. For example, it is plausible that BMI would have a small effect on MI risk for low-BMI individuals and a large effect for high-BMI individuals, but this type of nonlinearity cannot be gleaned from summary statistics (unless MI summary statistics were stratified by BMI). Eleventh, MR-style analyses have been applied to gene expression,^62–64^ and the potential for confounding due to pleiotropy in these studies could possibly motivate the use of LCV in this setting, but LCV is not applicable to molecular traits, which may be insufficiently polygenic for the LCV random-effects model to be well-powered. Finally, we have not exhaustively benchmarked LCV against every published MR method, but have restricted our simulations to the most widely used MR methods.^5, 7–11^ We note that there exist additional methods that aim to improve robustness by excluding or effectively down-weighting variants whose causal effect estimates appear to be outliers,^6, 12^ conceptually similar to the weighted median^8^ and mode-based estimator;^10^ however, we believe that any method that relies on genome-wide significant SNPs for a single one trait is likely to be confounded by genetic correlations (Figure 2). We further note that MR should ideally be applied to carefully curated sets of genetic variants that aim to exclude pleiotropic effects (MR with curation), but that curated sets of genetic variants are unavailable for most complex traits; in particular, it is difficult to compare LCV to MR with curation, as the performance of MR with curation will strongly depend on the quality of information used for curation, which can vary in practice.

Despite these limitations, for most pairs of complex traits we recommend using LCV instead of MR. When the exposure is a complex trait, MR is likely to be confounded by genetic correlations, and it may be impossible to identify valid instruments. However, there are several scenarios in which MR should be used, either in addition to or instead of LCV. First, when it is possible to produce a curated set of variants that are likely to represent valid instruments because they have a mechanistically direct effect on the exposure, it is appropriate to perform MR. For example, an MR analysis identified a causal effect of vitamin D on multiple sclerosis, utilizing genetic variants near genes with well-characterized effects on vitamin D synthesis, metabolism and transport; these variants all provided consistent estimates of the causal effect.^66^ As another example, cis-eQTLs can be used as genetic instruments to test for an effect of gene expression because they are unlikely to be confounded by processes mediated in trans, motivating applications of MR and related methods to gene expression^62–64^ (however, these studies also have other limitations, such as the high likelihood that GWAS SNPs may approximately colocalize with an eQTL^63, 65^). Second, when prior knowledge about likely pleiotropic factors is available, it is appropriate to perform MR in addition to LCV, either restricting to a curated set of variants without overt pleiotropic effects or correcting for these effects in a multivariate regression model.^4, 60^ Third, when one of the traits has low significance for nonzero heritability, LCV may produce unreliable estimates and MR should be used either instead of or in addition to LCV. Finally, well-powered MR studies can be used to show that two traits do not have a strong, fully genetically causal relationship, as confounding due to pleiotropy is more likely to lead to false positives than false negatives. In each case, MR should be performed with multiple genetic variants, a bidirectional analysis^11, 17^ should be performed to reduce the potential for confounding due to genetic correlations, and consistency of causal effect estimates across variants should be assessed both manually and analytically.^12^

## Acknowledgements

We are grateful to Ben Neale, Soumya Raychaudhuri, Chirag Patel, Sek Kathiresan, Bogdan Pasaniuc and Hilary Finucane for helpful discussions, and to Po-Ru Loh and Steven Gazal for producing BOLT-LMM summary statistics for UK Biobank traits. This research was conducted using the UK Biobank Resource under Application #16549 and funded by NIH grants R01 MH107649, U01 CA194393 and R01 MH101244.

## URLs

Open-source software implementing our method is available at github.com/lukejoconnor/LCV.

## Online Methods

### LCV model

The LCV random effects model assumes that the distribution of marginal effect sizes for the two traits can be written as the sum of two independent bivariate distributions (visualized in Figure 1c-e in orange and blue respectively): (1) a *shared genetic component*(*q*_1_*π, q*_2_*π*) whose values are proportional for both traits; and (2) an*even genetic component* (*γ*_1_, *γ*_2_) whose density is mirror symmetric across both axes. Distribution (1) resembles a line through the origin, and we interpret its effects as being mediated by a latent causal variable (*L*) (Figure 1a); distribution (2) does not contribute to the genetic correlation, and we interpret its effects as direct effects. Informally, the LCV model assumes that any asymmetry in the shared genetic architecture arises from the action of a latent variable.

In detail, the LCV model assumes that there exist scalars *q*_1_, *q*_2_, and a distribution (*π*, γ_1_, *γ*_2_)such that

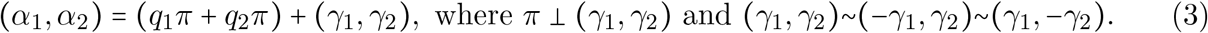

Here *α_k_* is the random marginal effect of a SNP of trait *k*, *π* interpreted as the marginal effect of a SNP on *L*, and*γ_k_* is interpreted as the non-mediated effect of a SNP on trait *k*. *α* and *π* (but not γ) are normalized to have unit variance, and all random variables have zero mean. (The symbol “~” means “has the same distribution as.”) *q*_1_, *q*_2_ are the model parameters of primary interest, and we can relate them to the mixed fourth moments, which are observable (equation (2)). In particular, this implies that the model is identifiable (except when the excess kurtosis *κ_π_* = 0; see Supplementary Note). We do not expect that *_ˇ_* will be exactly zero for any real trait, but there will be lower power for traits with higher polygenicity. Note that we have avoided assuming a particular parametric distribution.

The LCV model assumptions are strictly weaker than the assumptions made by MR. Like LCV, a formulation of the MR assumptions is that the bivariate distribution of SNP effect sizes can be expressed in terms of two distributions. In particular, it assumes that the effect size distribution is a mixture of (1’) a distribution whose values are proportional for both traits (representing all SNPs that affect the exposure Y1) and (2’) a distribution with zero values for the exposure Y1 (representing SNPs that only affect the outcome Y2). These two distributions can be compared with distributions (1) and (2) above. Because (1’) is identical to (1) and (2’) is a special case of (2), the LCV model assumptions are strictly weaker than the MR assumptions (indeed, much weaker). We also note that the MR model is commonly illustrated with a non-genetic confounder affecting both traits. Our latent variable *L* is a genetic variable, and it is not analogous to the non-genetic confounder. Similar to MR, LCV is unaffected by nongenetic confounders (such a confounder may result in a phenotypic correlation that is unequal to the genetic correlation).

The genetic causality proportion (gcp) is defined as:

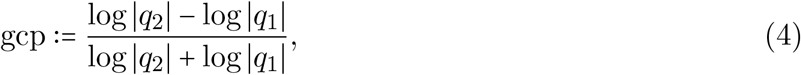

which satisfies

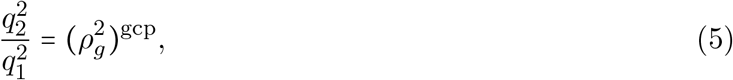

where the genetic correlation *ρ_g_* is equal to *q*_1_*q*_2_. gcp is positive when trait 1 is partially genetically causal for trait 2. When gcp = 1, trait 1 is fully genetically causal for trait 2: *q*_1_ = 1 and the causal effect size is *q*_2_ = *ρ_g_* (Figure 1b,e). The LCV model is broadly related to dimension reduction techniques such as Factor Analysis^67^ and Independent Components Analysis,^68^ although it differs in its modeling assumptions as well as its goal (causal inference); our inference strategy (mixed fourth moments) also differs.

Under the LCV model assumptions, we derive equation (2) as follows:

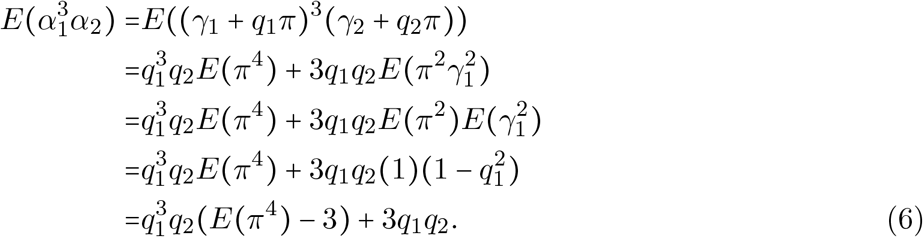

In the second line, we used the independence assumption to discard cross-terms of the form *γ_p_π*^3^ and 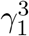*π*, and we used the symmetry assumption to discard terms of the form *γ*_1_ 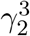. In the third and fourth lines, we used the independence assumption, which implies that *E* (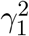*π*^2^) = *E*(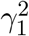)*E*(*π*^2^) = *E*(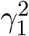) = 1 – 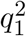. The factor *E*(*π*^4^) – 3 is the excess kurtosis of *π*, which is zero when *π* follows a Gaussian distribution; in order for equation (2) to be useful for inference, *E*(*π*^4^) – 3 must be nonzero, and in order for the model to be identifiable, *π* must be non-Gaussian (see Supplementary Note).

### Estimation under the LCV model

In order to estimate the gcp and to test for partial causality, we utilize six steps. First, we use LD score regression^19^ to estimate the heritability of each trait; these estimates are used to normalize the summary statistics. Second, we apply cross-trait LD score regression^16^ to estimate the genetic correlation; the intercept in this regression is also used to correct for possible sample overlap when estimating the mixed fourth moments. Third, we estimate the mixed fourth moments *E*(*α*_1_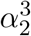) and *E* 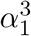*α*_2_) of the bivariate effect size distribution. Fourth, we compute test statistics for each possible of the gcp, based on the estimated genetic correlation and on the estimated mixed fourth moments. Fifth, we jackknife on these test statistics to estimate their standard errors, similar to ref. 19, obtaining a likelihood function for the gcp. Sixth, we obtain posterior means and standard errors for the gcp using this likelihood function and a uniform prior distribution. These steps are detailed below.

First, we apply LD score regression to normalize the test statistics. Under the LCV model, the marginal effect sizes for each trait, *α*_1_ and *α*_2_, have unit variance. We use a slightly modified version of LD score regression,^19^ with LD scores computed from UK10K data.^58^ In particular, we run LD score regression using a slightly different weighting scheme, matching the weighting scheme in our mixed fourth moment estimators; the weight of SNP *i* was:

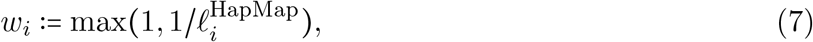

where 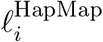 was the estimated LD score between SNP *i* and other HapMap3 SNPs (this is approximately the set of SNPs that were used in the regression). This weighting scheme is motivated by the fact that SNPs with high LD to other regression SNPs will be over-counted in the regression (see ref. 19). Similar to ref. 16, we improve power by excluding large-effect variants when computing the LD score intercept; for this study, we chose to exclude variants with *χ*^2^ statistic 30× the mean (but these variants are not excluded when computing 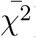). Then, we divide the summary statistics by *s* = 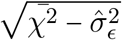, where 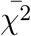 is the weighted mean *χ*^2^ statistic and 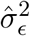 is the LD score intercept. (We also divide the LD score intercept by *s*^2^.) We assess the significance of the heritability by performing a block jackknife on *s*, defining the significance *Z_h_* as *s* divided by its estimated standard error.

Second, to estimate the genetic correlation, we apply cross-trait LD score regression.^16^ Similar to above, we use a slightly modified weighting scheme (equation (7)), and we exclude large-effect variants when computing the cross-trait LD score intercept. We assess the significance of the genetic correlation using a block jackknife.

Third, we estimate the mixed fourth moments *E*(*α*_1_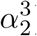 using the following estimation equation:

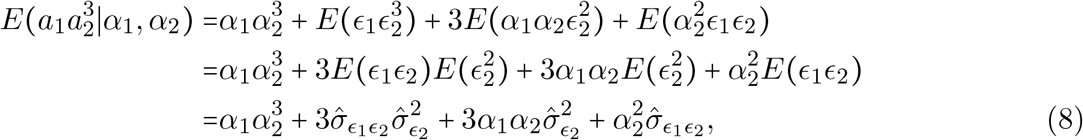

where *E*(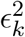) is the LD score regression intercept for trait *k* and 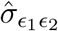 is the cross-trait LD score regression intercept. For simulations with no LD, we use *E*(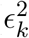) = 1/*sN_k_* and *E*(*∈*_1_*∈*_2_) = 0 instead of estimating these values.

Fourth, we define a collection of statistics *S*(*x*) for *x* ∈ *X* = {−1, −.01, −.02, *…*, 1} (corresponding to possible values of gcp):

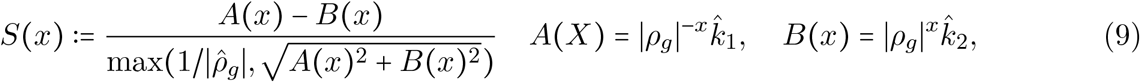

The motivation for utilizing the normalization by 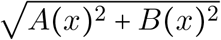 is that the magnitude of *A*(*x*) and *B*(*x*) tend to be highly correlated, leading to increased standard errors if we only use the numerator of *S*. However, the denominator tends to zero when the genetic correlation is zero, leading to instability in the test statistic and false positives. The use of the threshold leads to conservative, rather than inflated, standard errors when the genetic correlation is zero or nearly zero. We recommend only analyzing trait pairs with a significant genetic correlation, and this threshold usually has no effect on the results. It is also inadvisable to analyze trait pairs whose genetic correlation is non-significant because for positive LCV results, the genetic correlation provides critical information about the causal effect size and direction.

Fifth, we estimate the variance of *S(x*) using a block jackknife with *k* = 100 blocks of contiguous SNPs, resulting in minimal non-independence between blocks. Blocks are chosen to include the same number of SNPs, and the jackknife standard error is

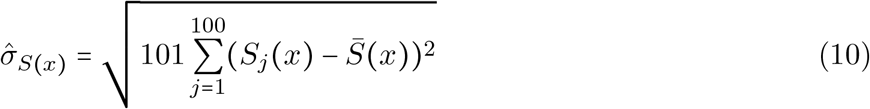

where *S_j_*(*x*) is the test statistic computed on blocks 1, …, *j* − 1, *j* + 1, *…*100 and *S*̅(*x*) is the mean of the jackknife estimates. We compute an approximate likelihood, *L*(*S*|gcp = *x*), by assuming (1) that *L*(*S*|gcp = *x*) = *L* (*S*|gcp = *x*) and (2) that if gcp = *x* then *S*(*x*)/*σ̂_S_(x*) follows a T distribution with 98 degrees of freedom.

Sixth, we impose a uniform prior on gcp, enabling us to obtain a posterior mean estimate of the gcp:

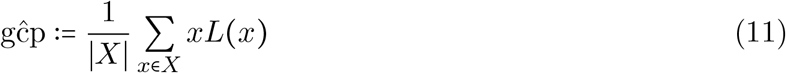

The estimated standard error is:

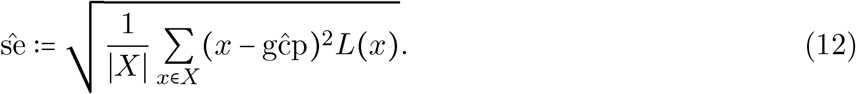

In order to compute p-values, we apply a T-test to the statistic *S*(0).

### Existing Mendelian randomization methods

#### Two-sample MR

As described in ref. 5, we ascertained significant SNPs (*p* < 5 × 10^−8^, *χ*^2^ test) for the exposure and performed an unweighted regression, with intercept fixed at zero, of the estimated effect sizes on the outcome with the estimated effect sizes on the exposure (in practice, a MAF-weighted and LD-adjusted regression is often used; in our simulations, all SNPs had equal MAF, and there was no LD). To assess the significance of the regression coefficient, we estimated the standard error as se = 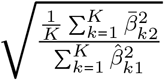 where *β̅*_*k*2_ is the *k*^th^ residual, *N*_2_ is the sample size in the outcome cohort, and *K* is the number of significant SNPs. This estimate of the standard error allows the residuals to be overdispersed compared with the error that is expected from the GWAS sample size. To obtain p values, we applied a two-tailed *t*-test to the regression coefficient divided by its standard error, with *K*− 1 degrees of freedom.

#### MR-Egger

As described in ref. 7, we ascertained significant SNPs for the exposure and coded them so that the alternative allele had a positive estimated effect on the exposure. We performed an unweighted regression with a fitted intercept of the estimated effect sizes on the outcome on the estimated effect sizes on the exposure. We assessed the significance of the regression using the same procedure as for two-sample MR, except that the *t*-test used *K*− 2 rather than *K* − 1 degrees of freedom.

#### Bidirectional MR

We implemented bidirectional mendelian randomization in a manner similar to ref. 11. Significant SNPs were ascertained for each trait. If the same SNP was significant for both traits, then it was assigned only to the trait where it ranked higher (if a SNP ranked equally high for both traits, it was excluded from both SNP sets). The Spearman correlations *r*_1_, *r*_2_ between the *z* scores for each trait was computed on each set of SNPs, and we applied a 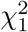 test to

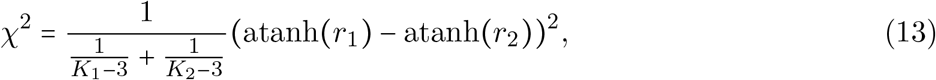

where *K_j_* is the number of significant SNPs for trait *j*. In ref. 11, the statistics atanh(*r_j_*) were also used, but a relative likelihood comparing several different models was reported instead of a p-value. We chose to report p-values for Bidirectional MR in order to allow a direct comparison with other methods.

#### Weighted median

As described in ref. 8, we ascertained significant SNPs for the exposure and computed ratio estimates and weights for each SNP. We computed the weighted median of the ratio estimates and estimated the standard error using a parametric bootstrap (100 bootstrap runs). We assessed significance using a Z test.

#### Mode based estimator

We ascertained significant SNPs for the exposure and computed ratio estimates for each SNP. We fit a curve to the observed ratio estimates using the Matlab fitdist() function with a bandwidth parameter as recommended in ref. 10, with uniform SNP weights. We verified that the Matlab fitdist() function produces identical curves as the original implementation in R. We computed the mode of the smoothed distribution and estimated its standard error using a parametric bootstrap (100 bootstrap runs). We assessed significance using a Z test.

#### Selection of genetic instruments on real data

For our applications of MR and related methods to real data, we selected genetic instruments using a greedy pruning procedure. We ranked all genome-wide significant SNPs for the exposure (*p* < 5 × 10^−8^) by *χ*^2^ statistic. Iteratively, we removed all SNPs within 1cM of the first SNP in the list, obtaining a set of independent lead SNPs separated by at least 1cM. We confirmed using an LD reference panel that our 1cM window was sufficient to minimize LD among the set of retained SNPs.

#### Application of MR to LDL and BMD

We applied two-sample MR (see above) to 8 curated SNPs that were previously used to show that LDL has a causal effect on CAD in ref. 3. 10 SNPs were used in ref. 3, of which summary statistics were available for 8 SNPs: rs646776, rs6511720, rs11206510, rs562338, rs6544713, rs7953249, rs10402271 and rs3846663.

## Simulations with no LD

In order to simulate summary statistics with no LD, first, we chose causal effect sizes for each SNP on each trait according to the LCV model. For all simulations except for Table S4, the causal effect size vector for trait *k* was

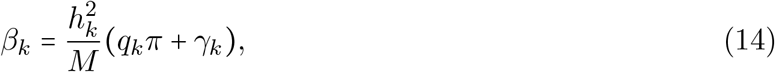

where in all simulations except for Table S5, *q_k_* was a scalar, and *π* and *γ_k_* were 1 × *M* vectors. In Table S5, *q_k_* was a 1 × 2 vector and *π* was a 2 × *M* matrix. Entries of *π* were drawn from i.i.d. point-normal distribution with mean zero, variance 1, and expected proportion of causal SNPs equal to *p_π_*. Entries of *γ_k_* were drawn from i.i.d. point-normal distributions with expected proportion of causal SNPs equal to *p_γk_*; we modeled colocalization between non-mediated effects by fixing some expected proportion of SNPs *p*_*γ*1_,_2_ < min(*p_γ_*_1_, *p_γ_*_2_) as having nonzero values of both *γ*_1_ and *γ*_2_. Then, we centered and re-scaled the nonzero entries of *π* and *γ_k_*, so that they had mean 0 and variance 1 and 1 − 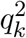, respectively.

For simulations in Table S4, effect sizes were drawn from a mixture of Normal distributions: there was a point mass at (0,0); a component with 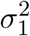 = 0, 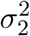 ≠ 0; a component with 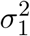 ≠ 0, 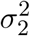 = 0; and a component with 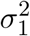 ≠ 0, 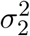 ≠_12_ = 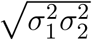. Values of *M, N_k_*, *N*_shared_, *ρ*_total_, *p_γk_*, *p*_*γ*1,2_, 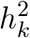, *p_π_*, *q_k_* for each simulation can be found in Table S1.

Second, we simulated summary statistics as

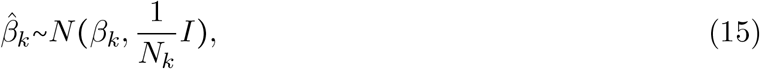

where *β_k_* is the vector of true causal effect sizes for trait *k* and *N_k_* is the sample size for trait *k*. When we ran LCV on these summary statistics, we used constrained-intercept LD score regression rather than variable-intercept LD score regression both to normalize the effect estimates^19^ and to estimate the genetic correlation,^16^ with LD scores equal to one for every SNP.

### Characterization of LCV model violations

In this subsection, we define partial genetic causality without making LCV (or other) model assumptions and characterize the type of LCV model violation that causes LCV to produce false positives and bias. There are two classes of LCV model violations: *independence violations* and *proportionality violations*. Roughly, independence violations involve a violation of the independence assumption between mediated effects (*π*) and direct effects (*γ*) while still satisfying a key proportionality condition related to the mixed fourth moments; as a result, independence violations are not expected to cause LCV to produce false positives (see Online Methods). Proportionality violations, on the other hand, violate this proportionality condition and are potentially more problematic. In order to make this characterization, it is necessary to define partial genetic causality in a more general setting, without assuming the LCV model. Partial genetic causality is defined in terms of the correlated genetic component of the bivariate SNP effect size distribution, which generalizes the shared genetic component modeled by LCV; unlike the shared genetic component, the correlated genetic component does not have proportional effects on both traits (but merely correlated effects).

#### Definition of partial genetic causality without LCV model assumptions

Let *A* = (*α*_1_, *α*_2_) be the bivariate distribution of marginal effect sizes, normalized to have zero mean and variance. First, we define an *even genetic component* of *A* as a distribution *T* = (*t*_1_, *t*_2_) that is independent of its complement *A* − *T* and that satisfies a mirror symmetry condition:

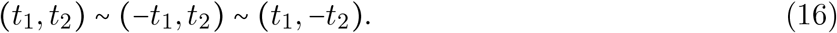

Equivalently, the density function of *T* is an even function of both variables. Note that an even genetic component does not contribute to the genetic correlation. In order to define the “correlated genetic component,” we would like to define a maximal even component, i.e. an even component that explains the largest possible amount of heritability for both traits. However, if *A* follows a Gaussian distribution, then there is no maximal even component: instead, the even genetic component that maximizes the proportion of trait 1 heritability explained fails to maximize the proportion of trait 2 heritability explained. This fact is related to the observation that the LCV model is non-identifiable when the effect size distribution for *L* follows a Gaussian distribution, and *only* when it follows a Gaussian distribution (see Supplementary Note). Generalizing this result, we conjecture that there exists an even component that is maximal up to a Gaussian term. More precisely, there exists a maximal even component *T*\* = (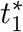, 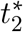) such that for any even component *T* = (*t*_1_, *t*_2_), there exists a (possibly degenerate) Gaussian random variable *Z* = (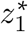, 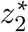) independent of *T*\* such that *T*\* + *Z* is an even component and *E*((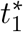 + *z*_1_)^2^) ≥ *E*(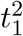) and *E*((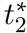 + *z*_2_)^2^) ≥ *E*(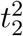).

We define the *correlated genetic component S* = (*s*_1_, *s*_2_) as the complement of the maximal even component and the Gaussian term. Trait 1 is defined as *partially genetically causal* for trait 2 if *E*(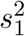) > *E*(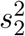), and vice versa. We may also define the genetic causality proportion using equation substituting *E*(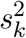) for 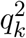 and *E*(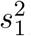)*E*(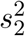) for 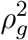. However, the interpretation of the gcp is not as clear in this more general setting. Note that the correlated genetic component may be identically 0, for example if *A* is bivariate Gaussian or if *A* itself is an even component; in both cases, there is no partial causality, and the genetic causality proportion is undefined. In practice, if the correlated genetic component is 0 or nearly 0, LCV will produce null p-values and low, noisy gcp estimates.

#### Independence violations and proportionality violations

The LCV model assumption is equivalent to the statement that the correlated genetic component resembles a line through the origin (and there is no Gaussian term): *S* = (*q*_1_*π, q*_2_*π*), for some random variable *π* and fixed parameters *q*_1_*q*_2_ such that *ρ_g_* = *q*_1_*q*_2_. Under the LCV model we refer to this distribution as the *shared genetic component* because its effects are fully shared (rather than merely correlated) between the two traits. This assumption enables an inference approach based on mixed fourth moments because it implies that the mixed fourth moments of the correlated component are proportional to the respective variances:

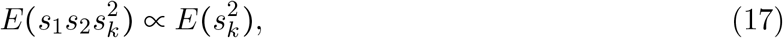

where under the LCV model, the proportionality constant is *q*_1_*q*_2_*E*(π^4^). However, the interpretation of the gcp is not as clear in this more general setting; in particular, a gcp of 1 implies that every SNP affecting trait 1 also affects trait 2, but not proportionally. Note that the correlated genetic component may be identically 0, for example if *A* is bivariate Gaussian or if *A* itself is an even genetic component; in both cases, there is no partial causality, and the genetic causality proportion is undefined. In practice, if the correlated genetic component is 0 or nearly 0, LCV will produce null p-values and low, noisy gcp estimates.

Intuitively, this type of violation arises as a result of non-independence between mediated effects (*π*) and direct effects (*γ*), causing “noise? from the direct effects to be incorporated into the correlated component. For this reason, we call such violations *independence violations*; genetic architectures that violate the proportionality condition we call *proportionality violations*. In the presence of an independence violation, we obtain the following moment condition, generalizing (6):

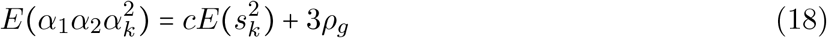

where *c* is a proportionality constant. In particular, if *E*(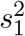) = *E*(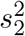) (no partial causality), then *E*(*α*_1_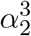) = *E*(*α*_2_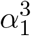), and LCV is expected to produce well-calibrated p-values. Conversely, under a proportionality violation, LCV is expected to produce inflated p-values under the null.

## Simulations with LD

In simulations with LD, we first simulated causal effect sizes for each trait in the same manner as simulations with no LD. Then, we obtained summary statistics in one of two ways, either using real genotypes or using real LD only.

For simulations with real genotypes modeling population stratification (Table 1g and Table S9), we chose effect sizes for each SNP and each trait from the LCV model with various parameters and multiplied these effect size vectors by real genotype vectors from UK Biobank,^27^ adding noise to obtain simulated phenotypes. For computational efficiency, we restricted these genotypes to chromosome 1 (*M* = 43k). We added stratification directly to the phenotype values along PC1 (computed on 43k SNPs and *N*_1_ + *N*_2_ individuals), with effect sizes 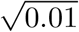 and 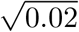 for trait 1 and trait 2, respectively. We then re-normalized phenotypes to have variance 1; afterwards, ~1% and ~2% of variance were explained by PC1 for each trait respectively. We estimated SNP effect sizes for each trait by correlating each SNP with the phenotypic values in *N_k_* individuals. In corrected simulations (Table S9b,d,f), we residualized the PC1 SNP loadings (computed on all *N*_1_ + *N*_2_ individuals) from the SNP effect estimates, a procedure which is effectively equivalent to correction of the individual-level data.^25^

For other simulations, we simulated summary statistics without first simulating phenotypic values, using the fact that the sampling distribution of *Z*-scores is approximately:^23^

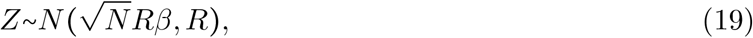

where *R* is the LD matrix and is the vector of true effect sizes. We estimated *R* from the *N* = 145*k* UK Biobank cohort using plink with an LD window size of 2Mb (*M* = 596*k*), which we converted into a block diagonal matrix with 1001 blocks. The number 1001 was chosen instead of the number 1000 so that the boundaries of these blocks would not align with the boundaries of our 100 jackknife blocks; the use of blocks allowed us to avoid diagonalizing a matrix of size 596k, while not significantly changing overall LD patterns (there are ~50, 000 independent SNPs in the genome, and 1001 << 50, 000). Because the use of a 2Mb window causes the estimated LD matrix to be non-positive semidefinite (even after converting it into a block diagonal matrix), each block was converted into a positive semidefinite matrix by diagonalizing it and removing its negative eigenvalues: that is, we replaced each block *A* = *V*Σ*V^T^* with the matrix *B*, where *B* = *V* max(0, Σ)*V^T^*. Then, because the removal of negative eigenvalues causes *B*′ to have entries slightly different from one, we re-normalized each block: *C* = *D*^−1/2^ *BD*^−1/2^, where *D* is the diagonal matrix corresponding to the diagonal of *B*. Even though the diagonal elements of *B* are close to 1 (mostly between 0.99 and 1.01), this step is important to obtain reliable heritability estimates using LD score regression because otherwise the diagonal elements of the LD matrix will be strongly correlated with the LD scores (*r*^2^ ≈ 0.5) and the heritability estimates will be upwardly biased, especially at low sample sizes.

We concatenated the blocks *C*_1_, …, *C*_1001_ to obtain a positive semi-definite block-diagonal matrix *R*′. We also computed and concatenated the matrix square root of each block. In order to obtain samples from a Normal distribution with mean *R*′ *β* and variance 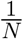 *R*′, we multiplied a vector having independent standard normal entries by the matrix square root of *R*′ and added this noise vector to the vector of true marginal effect sizes, *R*′ *β*. We computed LD scores directly from *R*. For simulations with sample overlap, the summary statistics were correlated between the two GWAS: the correlation between the noise term in the estimated effect of SNP *i* on trait 1 and the estimated effect of SNP *j* on trait 2 was *R*′_*ij*_*ρ*_total_*N*_shared_/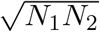, which is the amount of correlation that would be expected if the total (genetic plus environmental) correlation between the traits is *ρ*_total._^16^

### Outlier removal

In a secondary analysis, we applied an outlier removal procedure to determine whether our results on real traits using LCV were unduly influenced by individual loci. We computed the LCV test statistic *S*(0) (9) for each of the 100 jackknife blocks, discarded jackknife blocks that were ¿20 standard deviations from the mean, and re-ran the procedure iteratively until no outliers remained. For most trait pairs, this process results in the removal of 0 blocks; for a handful of trait pairs, it results in the removal of one or a few.

We do not recommend the broad use of this procedure, because outlier loci may contain valuable information. In particular, if any SNP affects trait 1 without affecting trait 2 proportionally, this suggests that trait 1 is not causal for trait 2. An alternative explanation is that its effect on trait 2 is masked by an opposing pleiotropic effect, either of the same causal SNP or of a different causal SNP at the same locus. If an outlier locus is to be removed, we recommend manually examining it and determining whether its removal can be justified or whether it provides competing statistical evidence against a causal effect.

